# An algal symbiont (*Breviolum psygmophilum*) responds more strongly to chronic high temperatures than its facultatively symbiotic coral host (*Astrangia poculata*)

**DOI:** 10.1101/2021.02.08.430325

**Authors:** Andrea N. Chan, Luis A. González-Guerrero, Roberto Iglesias-Prieto, Elizabeth M. Burmester, Randi D. Rotjan, John R. Finnerty, Iliana B. Baums

**Affiliations:** Biology Department, Pennsylvania State University, 208 Mueller Lab, University Park, PA 16801; Biology Department, Boston University, Boston, MA 02215

**Keywords:** coral, heat stress, climate change, facultative symbiosis, gene expression, photosynthesis, respiration, immune response

## Abstract

Scleractinian corals form the foundation of coral reefs by secreting skeletons of calcium carbonate. Their intracellular algal symbionts (Symbiodiniaceae) translocate a large proportion of photosynthate to the coral host, which is required to maintain high rates of calcification. Global warming is causing dissociation of coral host and algal symbiont, visibly presented as coral bleaching. Despite decades of study, the precise mechanisms of coral bleaching remain unknown. Separating the thermal stress response of the coral from the algal symbiont is key to understanding bleaching in tropical corals. The facultatively symbiotic northern star coral, *Astrangia poculata*, naturally occurs as both symbiotic and aposymbiotic (lacking algal symbionts) polyps – sometimes on the same coral colony. Thus, it is possible to separate the heat stress response of the coral host alone from the coral in symbiosis with its symbiont *Breviolum psygmophilum*. Using replicate symbiotic and aposymbiotic ramets of *A. poculata*, we conducted a chronic heat stress experiment to increase our understanding of the cellular mechanisms resulting in coral bleaching. Sustained high temperature stress resulted in photosynthetic dysfunction in *B. psygmophilum*, including a decline in maximum photosynthesis rate, maximum photochemical efficiency, and the absorbance peak of chlorophyll *a*. Interestingly, the metabolic rates of symbiotic and aposymbiotic corals were differentially impacted. RNAseq analysis revealed more differentially expressed genes between heat-stressed and control aposymbiotic colonies than heat-stressed and control symbiotic colonies. Notably, aposymbiotic colonies increased the expression of inflammation-associated genes such as nitric oxide synthases. Unexpectedly, the largest transcriptional response was observed between heat-stressed and control *B. psygmophilum*, including genes involved in photosynthesis, response to oxidative stress, and meiosis. Thus, it appears that the algal symbiont suppresses the immune response of the host, potentially increasing the vulnerability of the host to pathogens. The *A. poculata*-*B. psygmophilum* symbiosis provides a tractable model system for investigating thermal stress and immune challenge in scleractinian corals.

## Introduction

Scleractinian corals are foundation species on reefs because they secrete calcium carbonate at a high rate and so support the physical reef framework that provides habitat for a myriad of other taxa. Corals are able to maintain this energetically expensive biological process by forming an obligate symbiosis with dinoflagellates in the family Symbiodiniaceae (LaJeunesse *et al*., 2018, Odum & Odum, 1955). The algal symbionts translocate a significant portion of metabolic carbon as well as oxygen to the coral host, while the coral provides nutrients, protection from predators, and a stable location in the water column (Muscatine & Porter, 1977, Trench, 1979). This mutualism likely evolved during the mid-Triassic (Trench, 1997) and involves many physiological processes that are tightly linked between host and symbiont.

With the onset of anthropogenic climate change, sea surface temperatures are predicted to rise over the next century (Stocker *et al*., 2013). When temperatures exceed normal summer maximum temperatures globally, mass coral bleaching occurs (Hoegh-Guldberg, 1999). Coral bleaching is the breakdown of symbiosis (dysbiosis) between the coral host and algal symbiont, resulting in expulsion of the majority of algal cells (Weis *et al*., 2008). If temperatures do not return to normal, the corals often perish. However, it is difficult to separate the phenotypic response to elevated temperatures of the adult coral from its symbionts *in hospite*. Differentiating between the host’s and symbiont’s sensitivities to extreme temperatures is essential to inform management strategies and further our understanding of ecologically important associations between invertebrates and photosynthetic symbionts. Likewise, separating the host response from that of the symbiont is necessary to predict the potential for adaptation of existing symbioses to climate change.

Despite decades of study, the precise cellular mechanisms leading to coral bleaching are not fully understood (Davy *et al*., 2012). In order to separate the response of the cnidarian host from the response of the symbiont, previous studies have often utilized aposymbiotic coral larvae (Meyer *et al*., 2011, Polato *et al*., 2010, Rodriguez-Lanetty *et al*., 2009) or the cnidarian model sea anemone, Aiptasia (*Exaiptasia pallida*) (Dunn *et al*., 2007, Kitchen *et al*., 2017, Lehnert *et al*., 2014, Oakley *et al*., 2016, Weis *et al*., 2008). While the work comparing these symbiotic and aposymbiotic (lacking symbionts) anemones has been crucial to the current understanding of cnidarian-algal symbiosis, the energy budget of anemones and corals are not the same (due to the absence of a skeletal structure in *Aiptasia*) and thus their cellular responses to increased temperatures could differ markedly.

An emerging model system for cnidarian-algal symbiosis is the scleractinian temperate northern star coral, *Astrangia poculata* (Ellis & Solander, 1786, Peters *et al*., 1988). *A. poculata* is facultatively symbiotic with the dinoflagellate *Breviolum psygmophilum* (LaJeunesse *et al*., 2018, LaJeunesse *et al*., 2012), meaning it exists naturally as both symbiotic and aposymbiotic colonies. This coral uniquely possesses symbiotic (containing algal symbionts) and aposymbiotic (lacking algal symbionts) polyps on the same colony. This allows the researcher to manipulate the symbiotic state of the polyp while controlling for coral host genotype. An increasing number of studies in recent years have employed this model cnidarian to study the effects of ocean acidification and nutrients (Holcomb *et al*., 2012, Holcomb *et al*., 2010), wounding (Burmester *et al*., 2017, Burmester *et al*., 2018, DeFilippo *et al*., 2016), temperature (Burmester *et al*., 2017), and the associated microbiome (Sharp *et al*., 2017) on coral-algal symbiosis. Yet, to the authors’ knowledge, few studies have applied stressful high temperatures to study dysbiosis in *A. poculata* (Aichelman *et al*., 2019), and much work remains to be done.

For this study, we exposed genetically identical symbiotic and aposymbiotic fragments of *Astrangia poculata* to extreme temperatures to characterize dysbiosis, or the breakdown of symbiosis between *A. poculata* and *Breviolum psygmophilum*. While working with tropical corals often necessitates the application of acute heat shock techniques, we aimed to apply more realistic temperature ramping in our experiment while still applying thermal stress. Throughout our three-week experiment, we measured the stress response of *B. psygmophilum* at increasing temperatures using Pulse Amplitude Modulated (PAM) fluorometry, Photosynthesis-Irradiance curves, and reflectance spectra measurements. Likewise, we assessed the metabolic response of the coral host through respiration rate measurements in the absence of light. At the end of the experiment, RNA was extracted from all fragments and RNAseq analysis was conducted to distinguish between the coral host and symbiont responses to increased temperature stress.

## Materials and Methods

### Coral Colony Collection and Maintenance

Colonies of *Astrangia poculata* were collected at depths of 5.5-9.1 meters from Fort Wetherill State Park in Jamestown, Rhode Island in the fall of 2015. Large colonies (∼6cm in diameter) that were roughly half symbiotic and half aposymbiotic were targeted for collection. *A. poculata* colonies were temporarily maintained at the New England Aquarium prior to shipment to the Pennsylvania State University. Colonies were then acclimated in the Pennsylvania State aquarium system for approximately two weeks before fragmentation. Each colony was cut into roughly one symbiotic and one aposymbiotic portion using a diamond-coated dremel wheel. All colonies recovered for another two weeks before the second round of fragmentation, where we cut each symbiotic portion in half and each aposymbiotic portion in half. Thus, for each coral genet, we had four ramets, where two were symbiotic and two were aposymbiotic. This allowed us to control for host genetic variation with respect to symbiotic state while enabling the comparison of stress responses among host genotypes.

After fragmentation, all colonies of *A. poculata* were maintained in control conditions for approximately one year. Water temperature was maintained close to 18°C, salinity was maintained at 34 ppt, and water changes with artificial seawater (Instant Ocean Sea Salt) were conducted weekly. The aquarium system at Pennsylvania State University is closed, i.e. water is re-circulated through the coral tanks after being filtered in a sump system containing a filter sock (Innovative Marine AUQA Gadget 200 μm), protein skimmer (Vertex Omega 130), and bioballs. There are two independent sump reservoirs that each supply water to two paired aquarium tanks for a total of four tanks. Temperature can be independently regulated in the two systems (one sump tank plus two aquarium tanks).

### Experimental Design

Eight genets of *A. poculata*, each consisting of two symbiotic and two aposymbiotic ramets, were used in the temperature stress experiment (Figure 1). Ramets were randomly assigned to the “Control” or “High Temperature” treatment. Control ramets were maintained at 18°C throughout the three week experiment. Water temperatures in the experimental system were ramped every three days from a starting point of 18°C up until a final temperature of 30°C. Preliminary temperature stress experiments on *A. poculata* and the literature indicated that the upper thermal limit of *A. poculata* is higher than what individual colonies may experience in situ (Burmester *et al*., 2017, Jacques *et al*., 1983). Summer water temperatures in Naragansett Bay are usually between 18 and 24°C, however, previous work has reported that *A. poculata* colonies from this same region survive indefinitely in the laboratory at 27°C (Jacques *et al*., 1983). Thus, we chose a temperature that would be stressful for *A. poculata* to study the breakdown of symbiosis. During the experiment, the temperature was increased every three days. The experiment began with both systems at 18°C, during which baseline measurements were taken. On day three of the experiment, the temperature was raised 1°C every three hours until 22°C was reached. On day six, the temperature was again raised 1°C every three hours until 27°C was reached. Because these temperatures were not stressful for *A. poculata*, we only took measurements at 22°C and 27°C rather than at every 1°C increase. In Rhode Island, *A. poculata* colonies can experience diurnal temperature fluctuations greater than 5°C (Dimond & Carrington, 2007), and thus our ramping procedure was unlikely to be stressful. On day nine of the experiment (and every three days thereafter), the temperature was increased 1°C, until the final temperature of 30°C was reached on day 15. The temperature was maintained at 30°C for six additional days. All physiology measurements were taken during the two days between temperature changes to ensure that the corals had time to acclimate to each new temperature.

**Figure 1:**
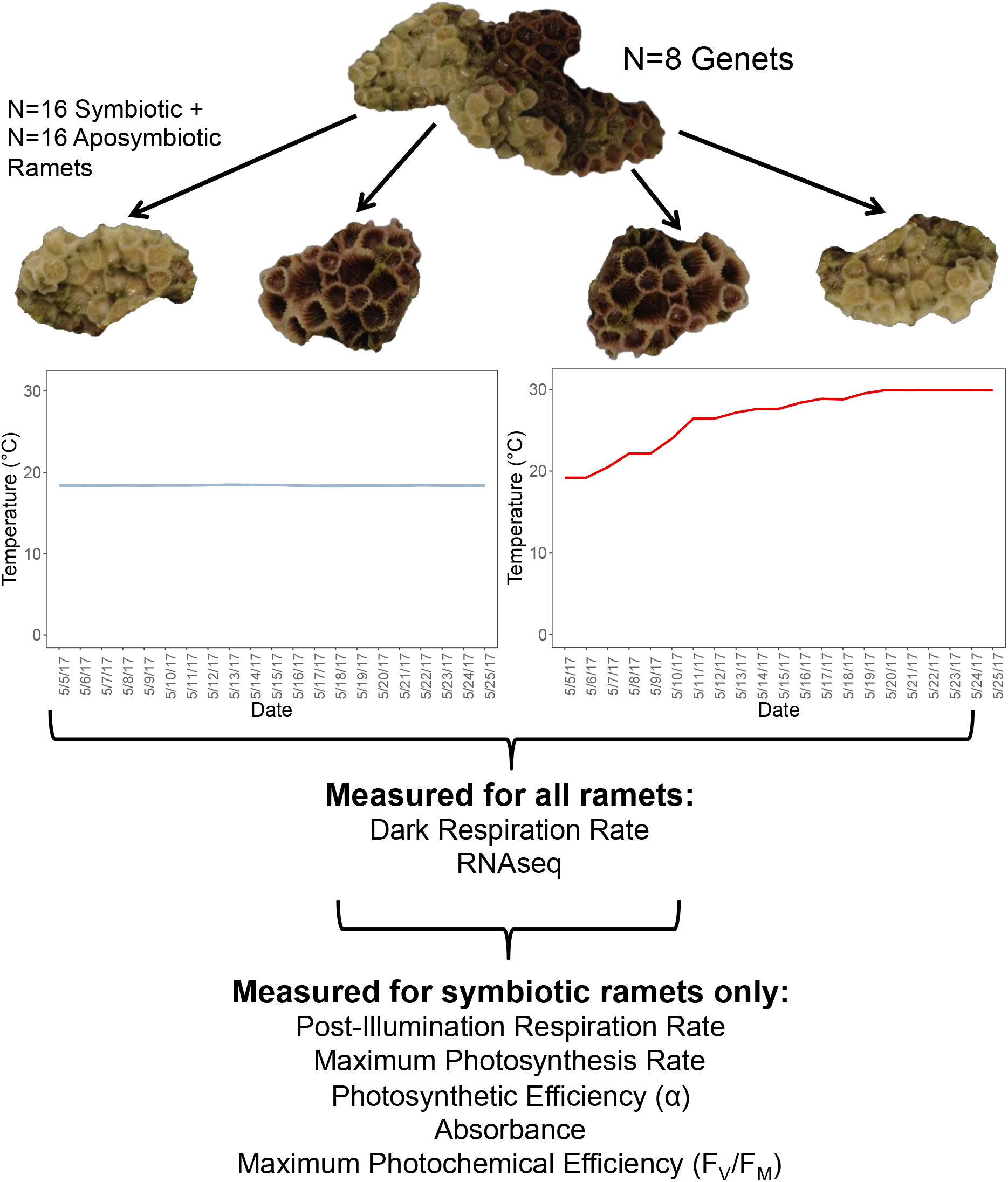
Overview of the heat stress experiment on paired symbiotic and aposymbiotic ramets of *Astrangia poculata*. Briefly, eight genets of *A. poculata* were divided into two symbiotic and two aposymbiotic ramets. One symbiotic and one aposymbiotic ramet from each genet were randomly assigned to the control (left graph) and experimental (right graph) treatments. The different measurements taken for all ramets and for only the symbiotic ramets are listed below the graphs.

All general system maintenance was conducted on days when no physiology measurements were taken, or every three days. The pH of the aquarium water in each system was measured using a Denver Instrument pH probe. Salinity was checked with a refractometer, and reverse osmosis-deionized water was added if the salinity was above 34ppt. Ammonia, nitrite, nitrate, and phosphate were measured using aquarium test kits (API Marine Saltwater Master Test Kit, Mars Fishcare, USA). In addition, the coral ramets were rotated to control for microcosm differences in light or water flow. A 19-liter (5 gallon) water change per system was conducted to remove nitrates from the water. Corals were fed freshly hatched brine shrimp nauplii every three days after photosynthesis and respiration measurements were taken. Illumination on a 12 hour light/12 hour dark cycle was provided by full spectrum LED lights dimmed to provide an average of 90 umol/m^2^s. Water temperature was recorded every 30 minutes using HOBO pendant loggers.

### Photosynthesis and Respiration Measurements

Photosynthesis and respiration measurements were taken every three days. Only half of the fragments could be measured in a single day due to the necessary incubation times, and so it took two days to complete one time point. The temperature was not ramped on the days that physiology measurements were taken. Coral fragments were incubated in custom-built hermetic chambers connected to a recirculating water bath that maintained the desired temperature in the chambers. The volume of filtered seawater used to fill each chamber was recorded for later normalization. Coral fragments were placed on a small stand within the chamber above a magnetic stir bar that thoroughly mixed the chamber water. Oxygen evolution was measured using an Optode System (FireSting, Germany). The incubations consisted of an initial measure of respiration in the dark (R_d_), nine increasing actinic light levels, and a final measure of the post-illumination respiration (R_l_). The light levels were previously calibrated in each chamber with a cosine light sensor of the diving PAM (Walz, Germany). For the aposymbiotic fragments, only dark respiration was measured in order to prevent *B. psygmophilum* infection in the light (Holcomb *et al*., 2012). Each coral fragment was incubated for a total of 105 minutes per trial, and each fragment was measured a total of seven times. The photosynthetic parameters were calculated according to standard methods (Osinga *et al*., 2012). Briefly, photosynthesis-irradiance (PE) curves were constructed by plotting the rate of change in oxygen concentration and irradiance. From these curves, photosynthetic parameters were calculated. The dark respiration rate (R_d_) is the slope of the linear regression relating change in oxygen concentration to time in the dark phase of the incubation. Photosynthetic efficiency (α) is the slope of the first few points of the PE curve before the saturation level of O_2_ is reached. Maximum photosynthesis rate (*P*_max_) is determined from the average of the last two points in the PE curve after reaching O_2_ saturation. Finally, the post-illumination respiration rate (R_l_) was measured during the second dark phase after fragments had already been exposed to increasing light levels. Our colonies of *A. poculata* were host to commensal organisms such as endolythic algae and tube worms that contributed to our measurements of photosynthesis and respiration. Due to the small size of our *A. poculata* fragments, and the need to immediately preserve tissue for RNAseq analysis, we could not measure photosynthesis and respiration of the skeleton alone to control for commensals, as in previous work (Jacques *et al*., 1983).

All measurements were normalized to the chamber water volume and coral fragment surface area. The surface area of each *A. poculata* fragment was measured using photogrammetry after corals were preserved in RNAlater. Fragments were placed one by one on a rotating turntable while a camera (Olympus M.Zuiko Digital, 14-42 mm lens) mounted on a tripod was used to automatically take a photograph approximately once per second. Photos were then imported into the Autodesk ReCap Pro photogrammetry software, where 3D models were generated. The surface area of the colony was measured by removing all parts of the model without coral tissue.

### Absorbance Estimations

Absorbance values were derived from reflectance spectra measurements, following the technical considerations of a previous study (Vásquez-Elizondo *et al*., 2017). Measurements were taken for each set of fragments on the same day that photosynthesis and respiration incubations were conducted. Fragments were submerged in a black container filled with artificial seawater to prevent extraneous light reflection and illuminated evenly with a custom semi-spherical light source suspended directly above the coral. An optical fiber (Ocean Optics, USA) was positioned at a 45° angle pointed at a coral polyp. A bleached skeleton of *A. poculata* was used as a reference for maximum light reflectance. For each fragment, we took three replicate reflectance measurements over the course of five minutes. Because the morphology of the *A. poculata* fragments varied, for some fragments it was not possible to obtain three replicate measurements with the time constraint. However, we were able to make at least one measurement per fragment per time point.

Reflectance measurements were analyzed in accordance with established methods (Enríquez *et al*., 2005, Vásquez-Elizondo *et al*., 2017). Briefly, replicate reflectance measurements were averaged, if applicable. Values were normalized to the amount of light reflected at 750nm, which is the maximum reflectance. Reflectance values were then converted to absorbance values, and plots of absorbance and wavelength were created.

### Maximum Photochemical Efficiency (Fv/Fm)

Maximum photochemical efficiency (Fv/Fm) was measured every day of the experiment at the same time each day. Corals were dark acclimated for 15 minutes before applying the saturating light pulse. A piece of PVC pipe was placed around the coral fragment to block light from reaching the other corals in the tank. Measurements were taken with a Diving PAM (Walz, Germany) and a 2 mm diameter optical fiber enabling single coral polyps to be measured.

### Statistical Analysis of Physiology Measurements

Dark respiration results for the symbiotic and aposymbiotic ramets, as well as the results for the other physiology measurements on the symbiotic ramets alone (including post-illumination respiration, maximum photosynthesis rate, photosynthetic efficiency (α), and the maximum photochemical efficiency) were analyzed using linear mixed effects models in R. The lmer function in the R package lmerTest was used to fit each model, with the fixed effects of symbiotic state (if applicable) and temperature, and random effects of coral ramets nested within genets. The anova function was then used to compute the analysis of variance tables, and the Tukey’s posthoc comparisons were calculated using the emmeans and contrast functions in the R package emmeans. Assumptions of normality, homoscedasticity, and independence were verified graphically in R with plots of the histogram of residuals, and the scatterplot of model residuals and fitted values.

### RNA Extraction, Library Preparation, and Sequencing

At the end of the experiment, all fragments were preserved in RNA*later* Stabilization Solution (Qiagen, Venlo, The Netherlands) and stored at 4°C for four days before being moved to the -20°C freezer. Total RNA was extracted using the mirVana miRNA Isolation Kit (Invitrogen, MA, USA), with an initial bead-beating step for tissue disruption. Coral tissue was cored from the polyps using a sterile forceps and placed in a 2mL tube filled with 0.5mL of plastic beads and 900 μl of lysis buffer. The tubes were then placed in the bead beater for three replications of bead-beating at maximum speed for 30 seconds, and then one minute on ice. Tubes were centrifuged at 12,000 rcf for 30 seconds and the supernatant was transferred to a new 2mL tube. The steps for the “Organic Extraction” and the “Total RNA Isolation Procedure” portions of the mirVana protocol were then followed. RNA was eluted in 50μl of elution solution (provided with the extraction kit). To avoid RNA degradation, 1.5μl of the RNAse inhibitor RiboGuard (Lucigen, purchased separately) was added to each sample. RNA was quantified using a Nanodrop ND-1000 spectrophotometer (Thermo Scientific, MA, USA) and RNA quality was assessed by running all samples on an Agilent 2100 Bioanalyzer (Agilent Technologies, CA, USA) using the RNA 6000 Nano Kit. Successful RNA extractions were then DNAse treated using the DNA-free kit (Ambion) to remove any contaminating DNA.

cDNA library preparation was conducted at the Pennsylvania State University Genomics Core Facility using the Illumina TruSeq Stranded mRNA protocol. cDNA libraries were sequenced using three Illumina HiSeq 2500 Rapid runs with two lanes per run. All samples were barcoded and loaded in each lane to avoid a lane effect and sequenced using 100-nucleotide single read sequencing.

### Differential Expression Analysis

FASTQ files from the three separate HiSeq Rapid runs were concatenated, yielding one FASTQ file per sample. FastQC reports were generated for each sample to identify overrepresented sequences (Andrews, 2010), such as TruSeq adapters and polyadenylated tails. Adapters were trimmed using Cutadapt with a quality cutoff of 15 and a minimum read length of 50 nucleotides (Martin, 2011). The program Kallisto (Bray *et al*., 2016) was used to pseudoalign trimmed reads to a reference transcriptome for the coral host *Astrangia poculata* (Burmester, 2017), and symbiotic samples were pseudoaligned to the reference transcriptome for *Breviolum psygmophilum* in culture (Parkinson *et al*., 2016), using 100 bootstrap replicates per sample. Pseudoalignment significantly reduces computation time by quantifying transcript abundance without aligning each read to a specific place within the transcript. The R package Sleuth (Pimentel *et al*., 2017) was then used to build models incorporating the factors of genotype, symbiotic state, RNA integrity number (RIN), and temperature treatment, and perform the differential expression analysis. The number and percentage of pseudoaligned reads to the coral host transcriptome are shown in Table 1, while the number and percentage of pseudoaligned reads to the symbiont transcriptome are shown in Table 2. The likelihood ratio test was used to compare nested models and identify differentially expressed transcripts. Sleuth incorporates the bootstrap results from Kallisto, enabling the program to distinguish between real biological expression differences and technical variation. Normalization in sleuth is the same as the “size factors” normalization procedure in DESeq (Anders & Huber, 2010).

**Table 1:**
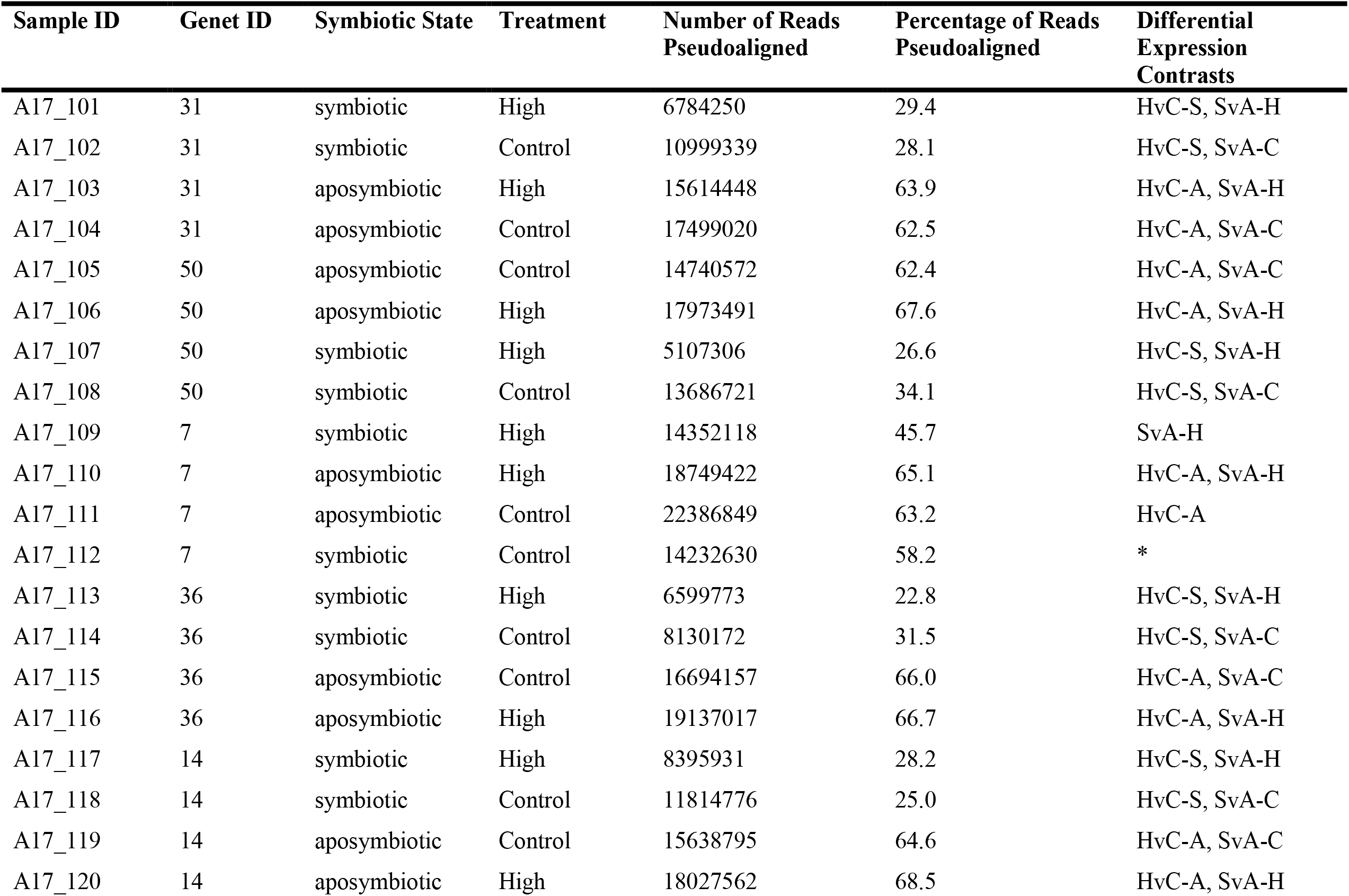
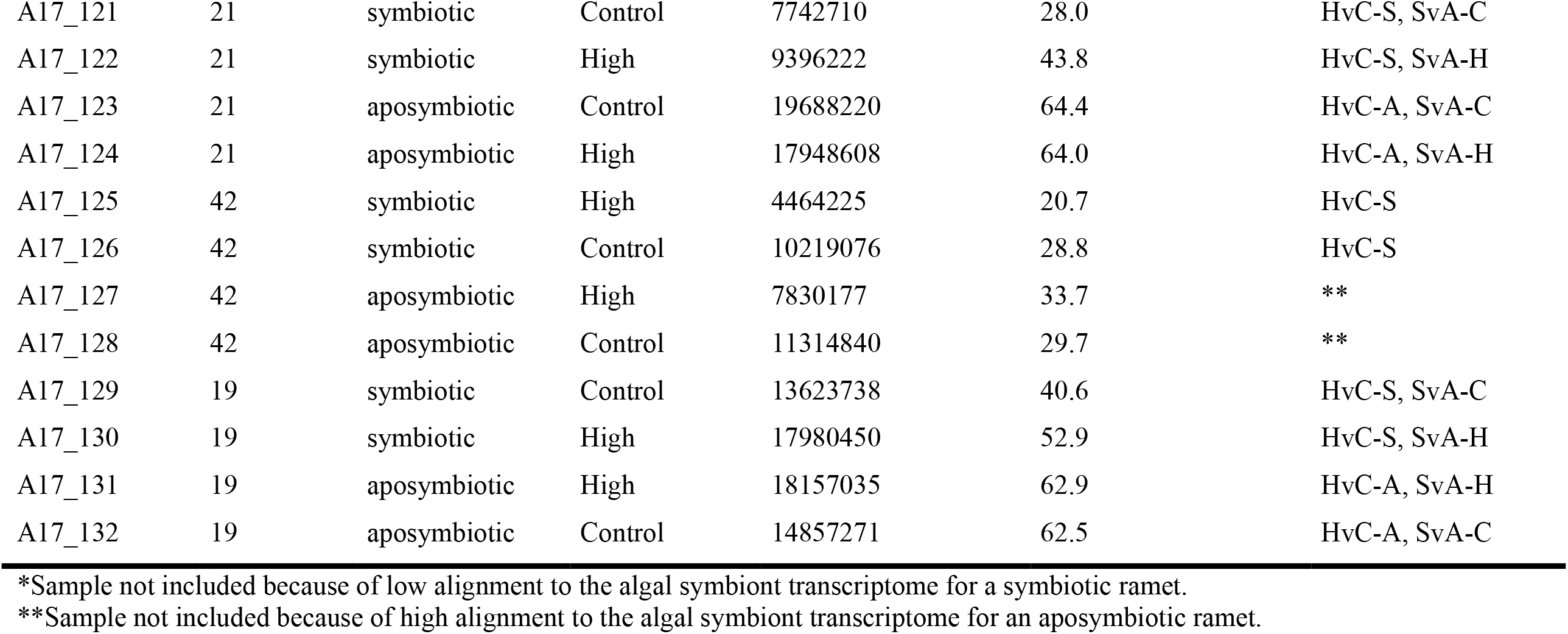
Samples included in differential expression contrasts for the coral host analyses. HvC-A: Heat vs. Control Contrast – Aposymbiotic Colonies Only; HvC-S: Heat vs. Control Contrast – Symbiotic Colonies Only; SvA-C: Symbiotic vs. Aposymbiotic Contrast – Control Colonies Only; SvA-H: Symbiotic vs. Aposymbiotic Contrast – Heat-Stressed Colonies Only

**Table 2:** Samples included in the Heat vs. Control Contrast for the algal symbiont differential expression analysis.

| Sample ID | Host Genet ID | Treatment | Number of Reads<br>Pseudoaligned | Percentage of Reads<br>Pseudoaligned |
| --- | --- | --- | --- | --- |
| A17_101 | 31 | High | 10446622 | 45.3 |
| A17_102 | 31 | Control | 19293038 | 49.2 |
| A17_107 | 50 | High | 9659785 | 50.4 |
| A17_108 | 50 | Control | 16710605 | 41.7 |
| A17_113 | 36 | High | 16243833 | 56.1 |
| A17_114 | 36 | Control | 11701965 | 45.4 |
| A17_117 | 14 | High | 14924576 | 50.1 |
| A17_118 | 14 | Control | 24291528 | 51.4 |
| A17_121 | 21 | Control | 13487662 | 48.8 |
| A17_122 | 21 | High | 5468062 | 25.5 |
| A17_125 | 42 | High | 10128722 | 47.0 |
| A17_126 | 42 | Control | 15771025 | 44.5 |
| A17_129 | 19 | Control | 7337254 | 21.9 |
| A17_130 | 19 | High | 4598122 | 13.5 |

### Functional Annotation of Differentially Expressed Transcripts

Gene names were assigned to differentially expressed transcripts by using BLASTx (National Center for Biotechnology Information) on the Pennsylvania State University Institute for CyberScience Advanced CyberInfrastructure high-performance computer cluster to compare transcript sequences against UniProt protein sequence databases (SwissProt and TREMBL) and the NCBI non-redundant (NR) protein database. The expect value (E value) cutoff was set to 10^−5^, which has been used previously in cnidarian genomic and transcriptomic studies. Putative genes were also assigned Kyoto Encyclopedia of Genes and Genomes (KEGG) orthology identifiers using the KEGG Automatic Annotation Server (v. 2.1), the single-directional best hit assignment method, and the representative gene set for eukaryotes, with the eight cnidarians (*Nematostella vectensis, Exaiptasia diaphana, Acropora digitifera, Acropora millepora, Pocillopora damicornis, Stylophora pistillata, Dendronephthya gigantea, Hydra vulgaris*) added (Moriya *et al*., 2007). The same settings were used to extract KEGG numbers for the *Breviolum psygmophilum* transcriptome (Parkinson *et al*., 2016), with the addition of the only dinoflagellate on the organism list (*Breviolum minutum*) to the template data set for the assignment of KEGG numbers. KEGG enrichment (overrepresentation) analysis was conducted using the *enrichKEGG* function and the KO database in the R package clusterProfiler (Yu *et al*., 2012). In order to improve visualization and interpretation of the enriched pathways, we constructed enrichment maps that cluster pathways sharing similar gene sets (Yu, 2018).

## Results

### Changes in Respiration and Photosynthesis Rates at Increasing Temperatures

The linear mixed effects model and ANOVA table were used to test for significant fixed effects of symbiosis and temperature, and significant random effects of coral ramets nested within genets on normalized respiration rates in the dark (R_d_, excluding the effects of photosynthesis). We found a significant effect of symbiotic state [F(1, 31.296)= 25.6168, *p*<0.0001], temperature [F(5, 179.389)= 41.9645, *p*<0.0001], and the interaction between symbiotic state and temperature [F(5, 179.389)= 2.4889, *p*=0.03303] on R_d_. Tukey post-hoc tests (α=0.05) were conducted to identify significant pairwise comparisons. As expected, at all temperatures, symbiotic ramets respired more than aposymbiotic ramets (Figure 2A), although this difference was not significant at 28 and 30°C. Little change in R_d_ was observed between 18°C and 27°C. Beyond 27°C, respiration rate increased for symbiotic and aposymbiotic ramets. Aposymbiotic ramets continued to increase their respiration rate at 30°C (Figure 2A), indicating that their enzymes and cellular processes maintained function at these high temperatures better than symbiotic ramets. Contrastingly, the symbiotic colonies decreased their respiration rates between 29 and 30°C – an indication of heat stress. As a result, R_d_ for symbiotic colonies did not differ significantly between 27°C and 30°C, whereas aposymbiotic colonies respired significantly more at the 30°C than at 27°C.

**Figure 2:**
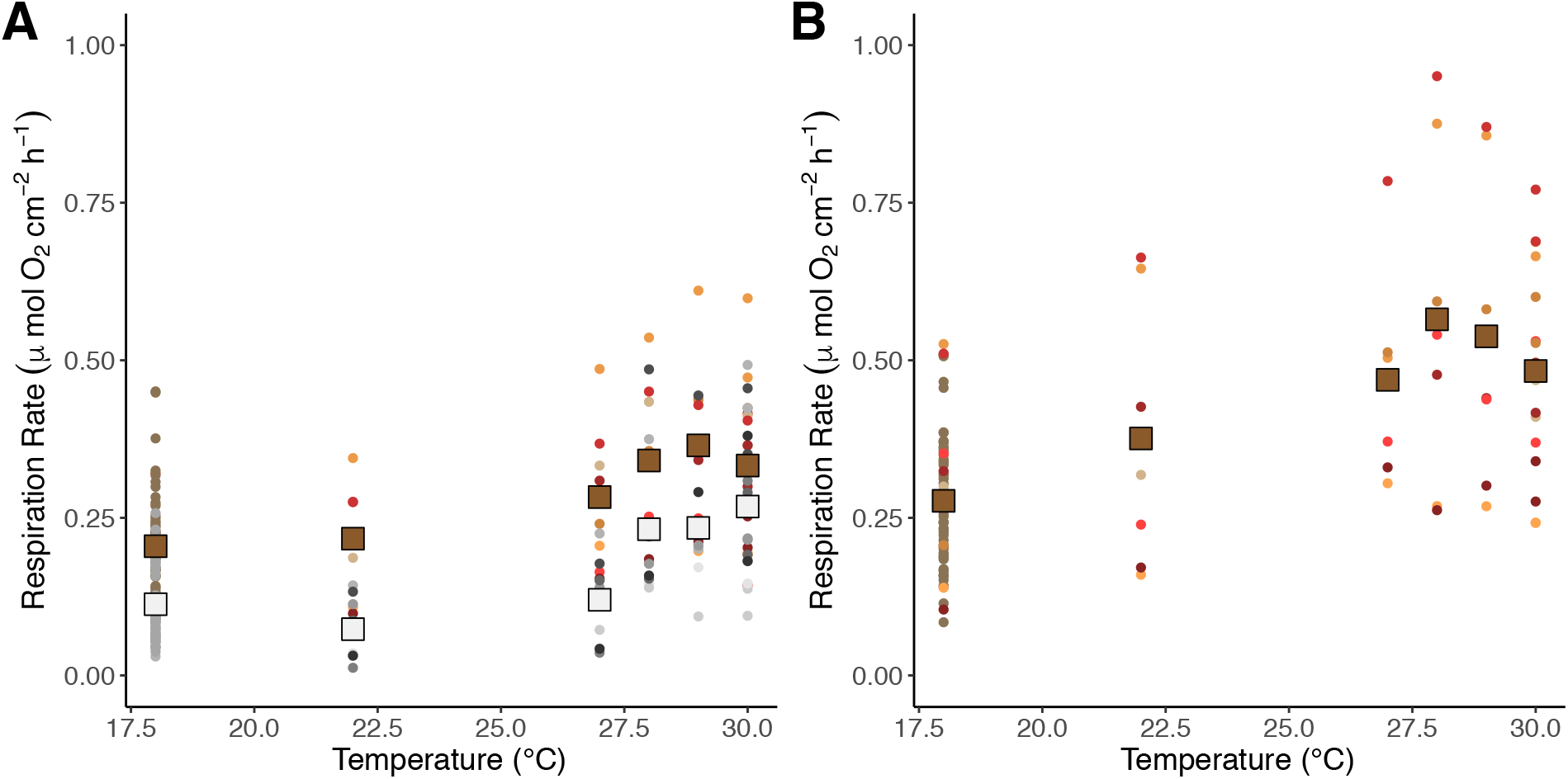
Graphs of mean dark respiration rates (R_d_) for aposymbiotic (pale gray squares) and symbiotic (brown squares) colonies (A), and mean post-illumination respiration rates (brown squares) for symbiotic colonies of *Astrangia poculata* (B) across temperatures. Point colors indicate measurements for the same ramets at each temperature level.

Post-illumination respiration (R_l_) was measured for the symbiotic ramets only. During post-illumination respiration, there is a high availability of oxygen and fixed carbon in the cells from symbiont photosynthesis. Because corals are oxyconformers, they will respire more in response to higher oxygen concentrations (Kühl *et al*., 1995, Rands *et al*., 1992). Thus, post-illumination respiration rates were higher than dark respiration rates at every measured temperature (Figure 2B). The Q_10_ temperature coefficient expresses the rate of change of a biological system because of increasing the temperature by 10°C. The Q_10_ values (calculated using respiration rates at 18°C and 28°C) for dark respiration and post-illumination respiration were 1.68 and 1.84, respectively. The lower Q_10_ value during dark respiration indicated that oxygen was somewhat limited under these conditions. A linear regression analysis between 18°C and 28°C yielded a significant positive relationship between temperature and post-illumination respiration (r^2^ = 0.208, *p* = 0.009). The ANOVA results revealed significant temperature effects on post-illumination respiration [F(5, 90.991) = 25.892, *p*<0.0001].

The average respiration rate was calculated by averaging dark and post-illumination respiration rates. Average respiration rate was used to calculate net photosynthesis rates from gross photosynthesis rates. The ANOVA revealed significant effects of temperature [F(5, 92.64) = 19.394, *p*<0.0001] on average net maximum photosynthesis rates (*P*_max_). With increasing temperature, *P*_max_ increased significantly from 18 to 27°C. *P*_max_ did not change significantly between 27 and 28°C, however, a significant decrease in *P*_max_ was observed between 28°C and 29°C as well as between 29°C and 30°C (Figure 3A, Tukey post-hoc tests).

**Figure 3:**
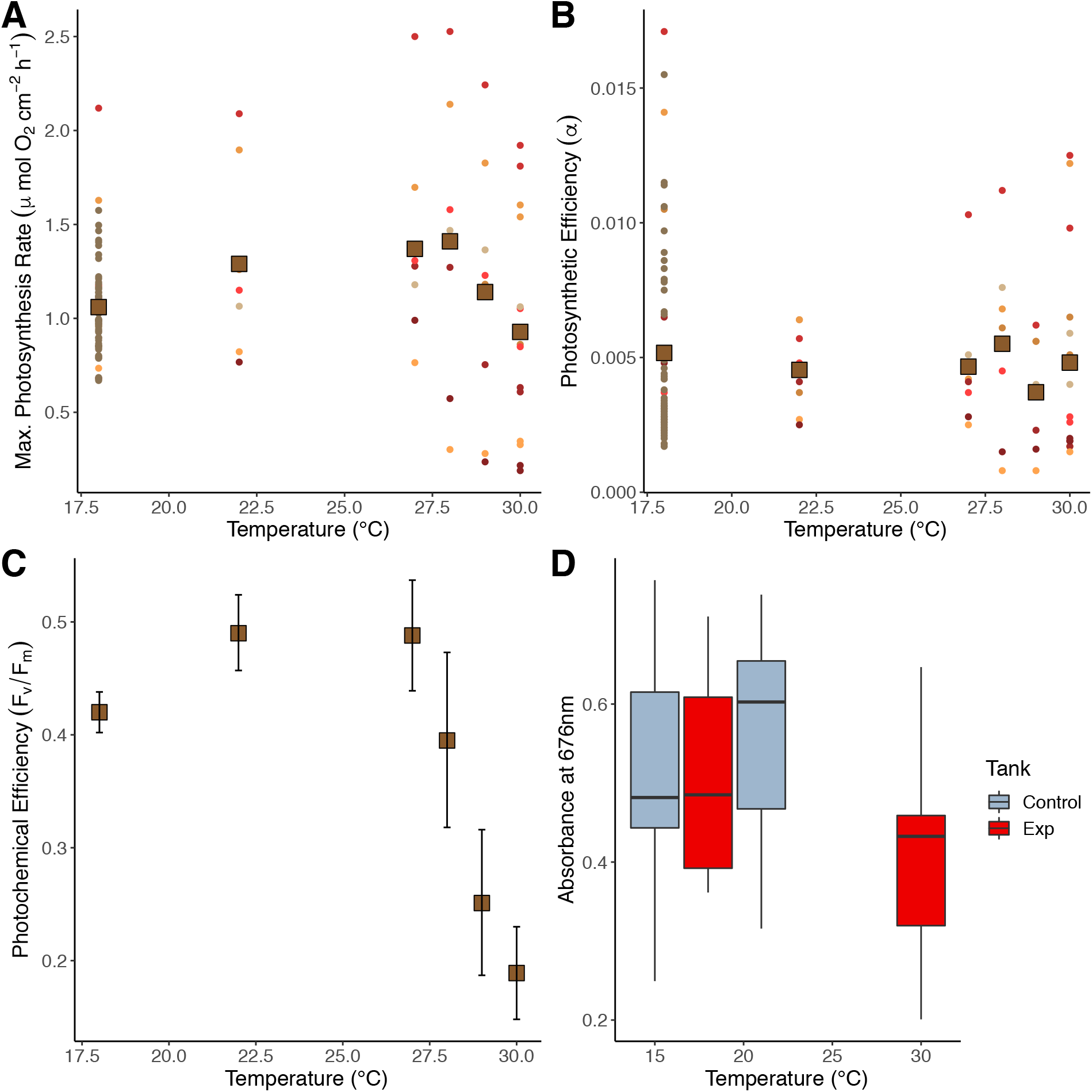
Average net maximum photosynthesis rate (*P*_max_, A) and photosynthetic efficiency (α, B) at increasing temperatures for symbiotic *Astrangia poculata* ramets. Point colors indicate measurements for the same ramets at each temperature level. The units for α are µmol O_2_ cm^-2^h^-1^ x (µmol quanta m^-2^s^-1^)^-1^. The average *P*_max_ and α at 18°C includes the control colonies as well as the colonies that were subjected to increasing temperatures. Normalized photochemical efficiency (quantum yield of photochemistry) for symbiotic *A. poculata* colonies with increasing temperatures (C). Averages and 95% confidence intervals are shown for each temperature level. Fv/Fm is the ratio of variable to maximum chlorophyll fluorescence. Chlorophyll *a* absorbance at 676 nm between control and experimental (Exp) symbiotic fragments of *Astrangia poculata* (D). Absorbance values for symbiotic ramets were calculated from reflectance spectra measured in accordance with standard procedures. The initial values when all ramets (control and experimental, n=8 each) were at 18°C did not differ significantly (t-test, *p*=0.903). However, a significant difference between absorbance values of the control (18°C) and experimental (Exp, 30°C) ramets was observed at the end of the experiment (t-test, *p*=0.0462).

Photosynthetic efficiency (α), or the initial slope of the Photosynthesis-Irradiance curve, was also calculated for each symbiotic coral ramet of *A. poculata* (Figure 3B). Notably, the residuals for this model were right-skewed and there were some patterns in the residual vs. fitted values plot. However, since linear mixed models are rather robust to some violations of their assumptions, we proceeded with the ANOVA. Throughout the temperature increase during the experiment, we found no significant change in photosynthetic efficiency [F(5, 74.21) = 1.6496, *p*=0.1575].

### Changes in Maximum Photochemical Efficiency (Fv/Fm) at Increasing Temperatures

The maximum photochemical efficiency (Fv/Fm) decreased sharply above 27°C for the symbiotic *A. poculata* ramets (Figure 3C). Temperature [F(5, 101.68)= 39.818, *p*<0.0001] significantly affected Fv/Fm (ANOVA results from linear mixed effects model). However, maximum photochemical efficiency was only significantly lower in symbiotic *A. poculata* ramets at 29°C and 30°C, compared to measurements at all lower temperatures (Tukey post-hoc tests). A linear regression analysis of the photochemical efficiency values between 27 and 30°C revealed a significant negative relationship with temperature (r^2^ = 0.431, *p* < 0.001).

## Absorbance estimations

We compared the difference in absorbance between the control *A. poculata* fragments and the heat-stressed *A. poculata* fragments, focusing on the absorbance at 676 nm corresponding to peak absorbance of chlorophyll *a*. The chlorophyll *a* peak was well defined in the combined absorbance spectra. Other photosynthetic pigments contributed to the absorbance spectra at different wavelengths such as carotenoids and chlorophyll *b* from the endolithic algae associated with the *A. poculata* skeleton. At the start of the experiment, the absorbance at 676 nm did not differ between the control and experimental coral fragments (t-test, *p*=0.903). However, after the experimental fragments were exposed to increasing temperatures for three weeks, the absorbance at 676 nm decreased significantly compared to the controls (Figure 3D, *p*=0.0462). Thus, while the experimental *A. poculata* fragments did not lose all their algal symbionts the way tropical corals do during a thermal stress event, these *A. poculata* fragments were developing a bleached phenotype at the end of our experiment.

### Differential Expression Contrasts

After removing samples with low mapping percentages to the reference transcriptomes, or aposymbiotic samples with unusually high symbiont reads (Table 1), overall differential expression patterns were compared using principle component analyses (PCA). The PCA showing the variation in gene expression for all samples mapped to the coral host transcriptome is shown in Figure 4A. As expected, samples clustered by temperature treatment along the first principle component (PC1 = 38.17%), regardless of symbiotic state. Figure 4B shows the PCA for just the symbiotic samples that were mapped to the *B. psygmophilum* transcriptome. The highest variability in gene expression for these samples was between symbiont strains within the high temperature treatment (PC1 = 86.43%). The second principle component separated samples by temperature treatment, but explained a much smaller percentage of the variation (PC2 = 8.34%).

**Figure 4:**
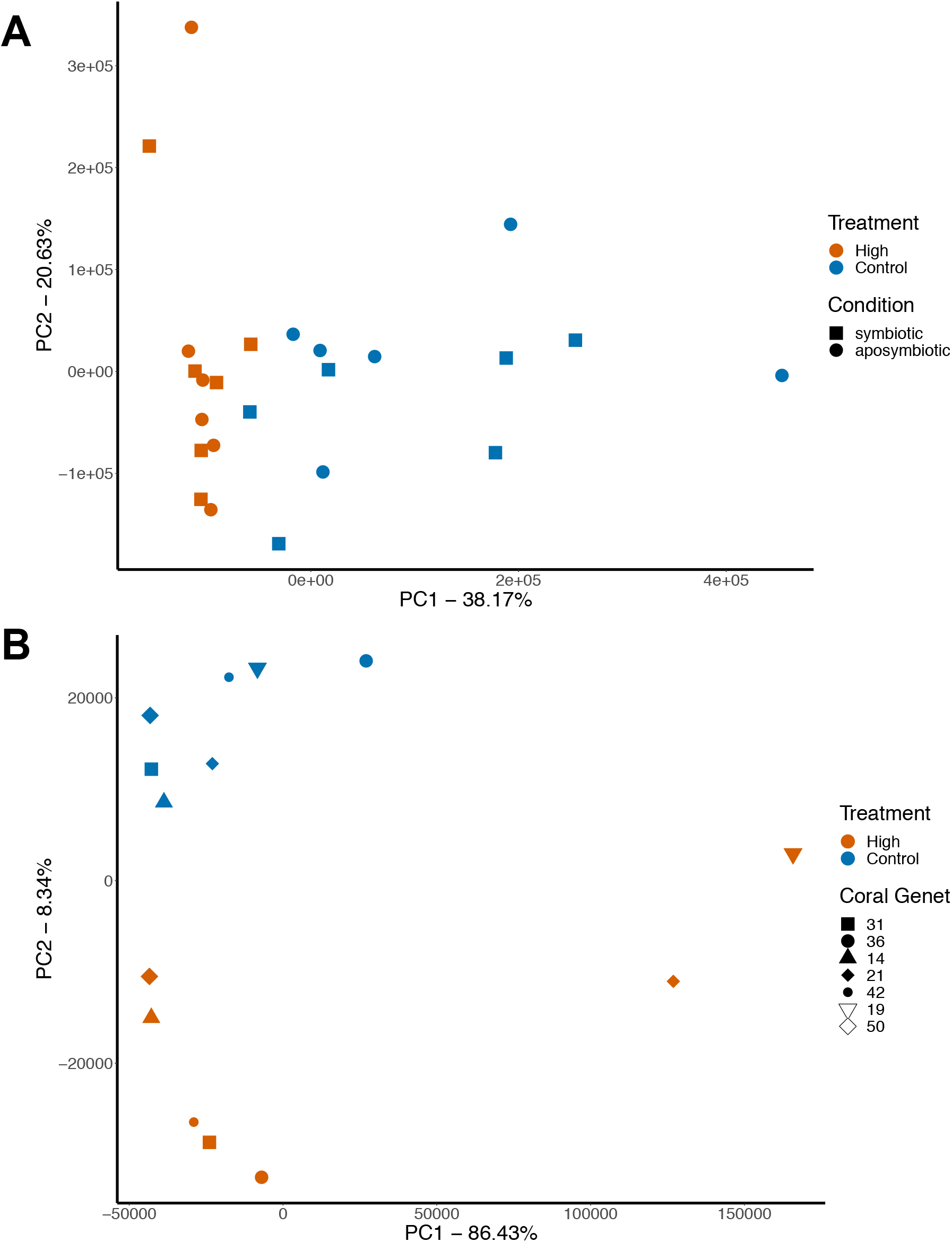
Principle component analyses of differential gene expression of the coral holobiont (A) and algal symbiont *in hospite* (B). The percentage of variation explained by the first two components is indicated on each axis.

### Heat vs. Control Contrast – Aposymbiotic Ramets Only

For the aposymbiotic coral colonies, there were 76 differentially expressed transcripts between the control and high temperature treatments (Figure 5A). Informative annotation was obtained for all but 18 of the 76 transcripts between the three databases (Swissprot, Trembl, and NCBI). Best-hit results were largely consistent between databases, with a majority of coral species as the source organisms for protein sequences from the Trembl and NCBI databases.

**Figure 5:**
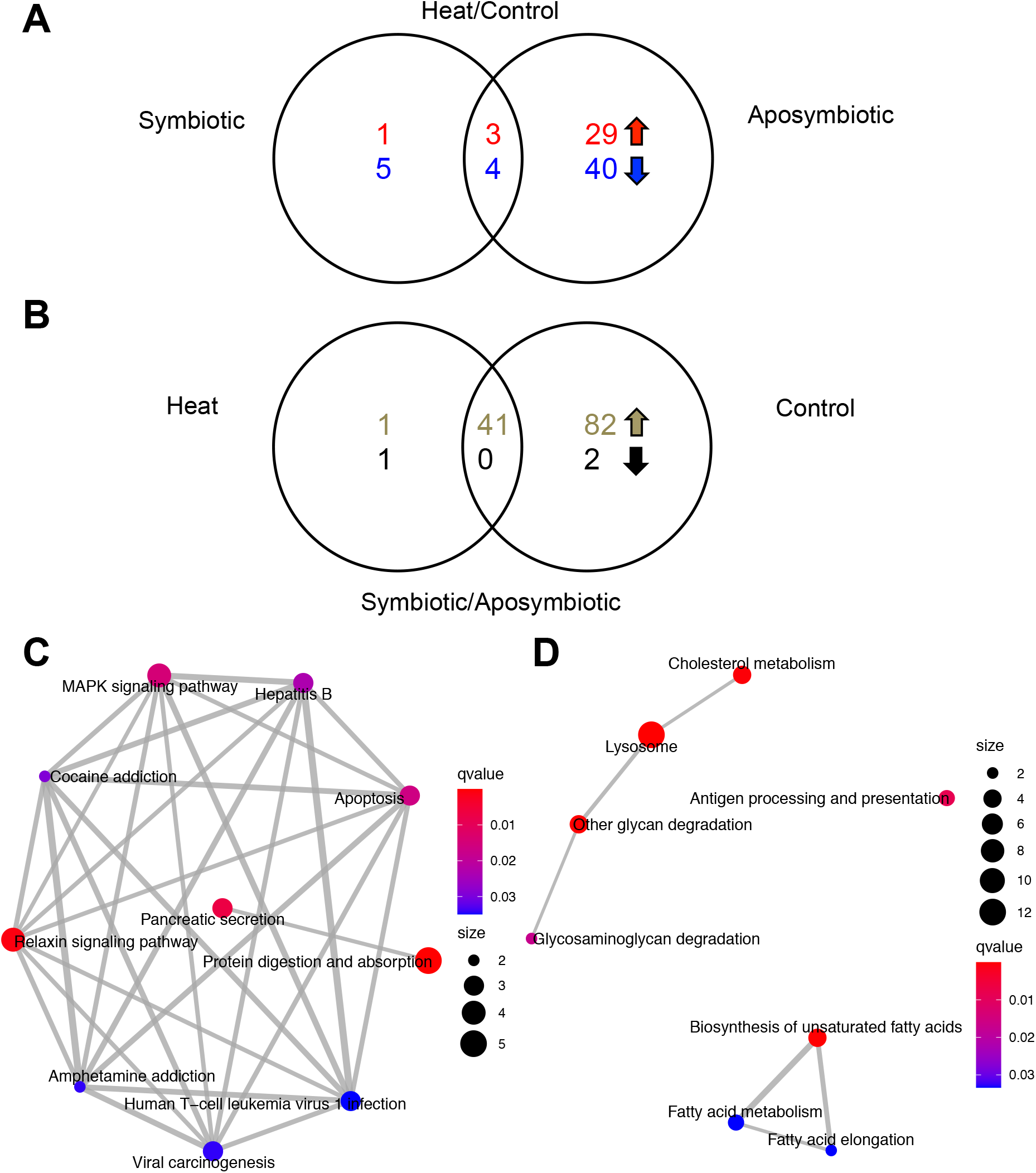
Venn diagrams showing the number of differentially expressed transcripts (DET) and enriched KEGG pathways in response to temperature stress for *Astrangia poculata*. (A) The number of DETs in symbiotic heat-stressed ramets relative to controls (left circle) and aposymbiotic heat-stressed ramets relative to controls (right circle). (B) The number of DETs in heat-stressed symbiotic ramets relative to aposymbiotic ramets (left circle) and control symbiotic ramets relative to aposymbiotic ramets (right circle). Top numbers indicate the number of upregulated transcripts while bottom numbers show the number of downregulated transcripts. Enrichment maps showing the ten significantly enriched KEGG pathways (circular nodes) when comparing heat-stressed and control aposymbiotic *A. poculata* (C) and the eight enriched pathways when comparing symbiotic and aposymbiotic control *A. poculata* (D). Gray lines connect pathways with overlapping genes. The color of each circle indicates the significance (q-value) and the size shows the number of genes in that pathway.

In the heat stressed aposymbiotic colonies, there were 32 and 44 upregulated and downregulated transcripts, respectively, relative to the control colonies. The most upregulated transcripts (measured by log_2_ fold change) included a tumor necrosis factor-related protein, nitric oxide synthases, and a member of the small heat shock protein family (*hsp20*) associated with chaperone-mediated protein folding. Downregulated genes relative to controls coded for enzymes involved in the breakdown of complex carbohydrates, a mitochondrial transferase, and a protein that catalyzes the elongation of very long-chain fatty acids.

KEGG enrichment analysis for the aposymbiotic heat vs. control contrast yielded ten enriched pathways that included between two and five genes each (Figure 5C). The most significant KEGG pathway was protein digestion and absorption with five genes that were all downregulated in heat-stressed colonies. Three of these genes coded for collagen alpha chains, and the remaining two were extracellular or secreted peptidases. The relaxin signaling pathway was also enriched and included a nitric oxide synthase that was highly upregulated in heat stressed corals, as well as two upregulated transcription factors, and a downregulated collagen alpha chain. Nitric oxide synthases produce the free radical signaling molecule nitric oxide (*NO*). This important signaling molecule is part of the invertebrate immune response (Ellis *et al*., 2011). In this pathway, the collagen plays a role in anti-fibrosis. The MAPK signaling pathway was also enriched, and included a highly upregulated molecular chaperone (part of the small heat shock protein family) associated with protein refolding in response to heat in the fruit fly, *Drosophila melanogaster* (Vos *et al*., 2016). Notably, the apoptosis pathway was also significantly enriched, but only included three genes. The most upregulated gene in this pathway was a cyclic AMP-dependent transcription factor that is important for preventing apoptosis.

### Heat vs. Control Contrast – Symbiotic Ramets Only

There were only 13 differentially expressed transcripts between the heat-stressed symbiotic and control symbiotic ramets (six exclusive to the symbiotic coral contrast, seven shared with the aposymbiotic contrast), all of which were annotated with informative protein descriptions from the Swiss-Prot curated database (Figure 5A). Generally, the source organisms for the Swiss-Prot protein sequences are from model systems such as mouse (*Mus musculus*), although similar annotations were obtained from the Trembl database, where the source organisms were all stony corals or other cnidarians. Three of these transcripts were upregulated in the high temperature treatment, while the other ten were downregulated. The heat stressed corals significantly upregulated two genes involved in transcription regulation. These included a serrate RNA effector molecule homolog involved in mRNA processing and a transcription factor involved in the negative regulation of gene expression and transcription. The third upregulated gene was cubilin, which helps cells uptake lipoproteins, vitamins, and iron for further metabolism. Downregulated genes included those involved in carbohydrate metabolism, protein folding, molecular transport, and fatty acid chain elongation. An influenza virus NS1A-binding protein homolog was another downregulated gene in heat stressed symbiotic colonies. This protein helps prevent cell death from destabilizing actin filaments in mouse models (Matsudo *et al*., 2006).

KEGG enrichment analysis was conducted to identify overrepresented pathways in our differentially expressed transcripts. Six pathways were significantly enriched, with only one or two genes per pathway. Because so few genes were identified per pathway, these were likely only identified as ‘enriched’ because of the small input of K numbers. Thus, no further pathway analysis was conducted for this set of differentially expressed genes.

### Symbiotic vs. Aposymbiotic Contrast – Control Ramets Only

Testing for differential expression between the control symbiotic and aposymbiotic ramets revealed 125 differentially expressed transcripts, all of which were upregulated in symbiotic colonies except two (Figure 5B). Of these 125 transcripts, 41 were shared with the heat-stressed symbiotic vs. aposymbiotic comparison (next section). Informative annotations were retrieved for 97 transcripts. The two genes downregulated in the control symbiotic colonies were a serine/threonine protein kinase and a skeletal organic matrix protein. In the control symbiotic colonies, the most upregulated genes with informative annotation included a mitochondrial ATP synthase, a serine/threonine-protein kinase, and cathepsin (aids protein turnover).

KEGG enrichment analysis yielded eight significant pathways (Figure 5D). The first pathway involved lysosome activity and included twelve genes upregulated in symbiotic corals. These included cathepsins (proteases), arylsulfatase B, cholesterol transporters, a glucosylceramidase (catalyzes a reaction that yields glucose and ceramide), a lipase, a peptidase, and a protein important for carbohydrate breakdown (mannosidase). The second pathway was for “Other glycan degradation”, which also included a glucosylceramidase and other enzymes involved in carbohydrate breakdown. Cholesterol metabolism was also enriched, and included four genes upregulated in symbiotic control corals. Cholesterol transporters, a lipase, and an adapter protein important for endocytosis were all part of the cholesterol metabolism pathway. Another notable enriched pathway was antigen processing and presentation, which included upregulated cathepsins and a lysosomal thiol reductase important for preparing antigens for presentation to immune cells.

### Symbiotic vs. Aposymbiotic Contrast – Heat-Stressed Ramets Only

An examination of differentially expressed transcripts between symbiotic and aposymbiotic colonies that were all heat stressed revealed 42 upregulated transcripts in symbiotic ramets and only one transcript downregulated in symbiotic ramets (Figure 5B). Forty-one of these transcripts were also differentially transcribed under control conditions between aposymbiotic and symbiotic ramets (previous section). These results show that heat stress reduced the differences in transcription between aposymbiotic and symbiotic states (from 125 differentially expressed transcripts to 43). Of the 43 differentially expressed transcripts, 15 yielded no annotation results from the three protein databases, or returned uninformative annotation (e.g. uncharacterized protein). The sole downregulated gene in symbiotic heat-stressed colonies was a dioxygenase that regulates actomyosin processes and is involved in demethylation of proteins and DNA.

Genes that were strongly upregulated in symbiotic heat-stressed colonies relative to aposymbiotic heat-stressed colonies included an arylsulfatase B involved in autophagy and response to stimuli, a small amino acid transporter, and a DNA repair enzyme that binds damaged DNA. Other notable upregulated genes include a mitochondrial ATP synthase, carbonic anhydrase 2, and a glutaredoxin arsenate reductase. All of these were also upregulated in symbiotic control colonies. KEGG enrichment analysis was conducted for this gene set, but no pathways were identified as enriched.

### Heat vs. Control Contrast for the Algal Symbiont

Breviolum psygmophilum Unexpectedly, the largest number of differentially expressed transcripts was found between heat stressed and control samples of the intracellular algal symbiont, *Breviolum psygmophilum*. Using the published *B. psygmophilum* transcriptome derived from cultured algal cells as the mapping reference, we found 11,514 differentially expressed transcripts. After calculating the log_2_ fold changes for the set of differentially expressed transcripts in *B. psygmophilum*, we found 5,736 transcripts upregulated in heat-stressed samples and 5,778 that were downregulated. Most of these transcripts had small fold changes, with only 318 transcripts up- or downregulated by a log_2_ fold change of 1.5 or more in either direction. The differentially expressed transcripts with the largest fold change increase in heat-stressed samples included a metal-binding acid phosphatase with affinities for iron and zinc, a sodium channel protein, and an ammonium transporter implicated in ammonium uptake. Genes with the largest fold changes that were significantly downregulated in heat-stressed samples included a retrovirus-related Pol polyprotein from transposon TNT 1-94 (implicated in heavy metal stress tolerance), and a triosephosphate isomerase involved in gluconeogenesis and glycolysis.

KEGG enrichment analysis on the entire set of *B. psygmophilum* differentially expressed transcripts (n=11,514) produced 58 significantly enriched KEGG pathways (Figure 6). The pathways with the smallest q-values (most significantly enriched) included “Spliceosome”, “Ribosome”, and “Protein processing in endoplasmic reticulum”, which are all involved in protein synthesis and processing. One of the largest networks of pathways included cell cycle and meiosis pathways, such as the “Oocyte meiosis” and “Meiosis - yeast” pathways (Figure 6). Another enrichment map cluster consisted of pathways involved in carbon metabolism, such as “Carbon fixation in photosynthetic organisms”. Smaller clusters of pathways corresponded to autophagy and DNA repair mechanisms.

**Figure 6:**
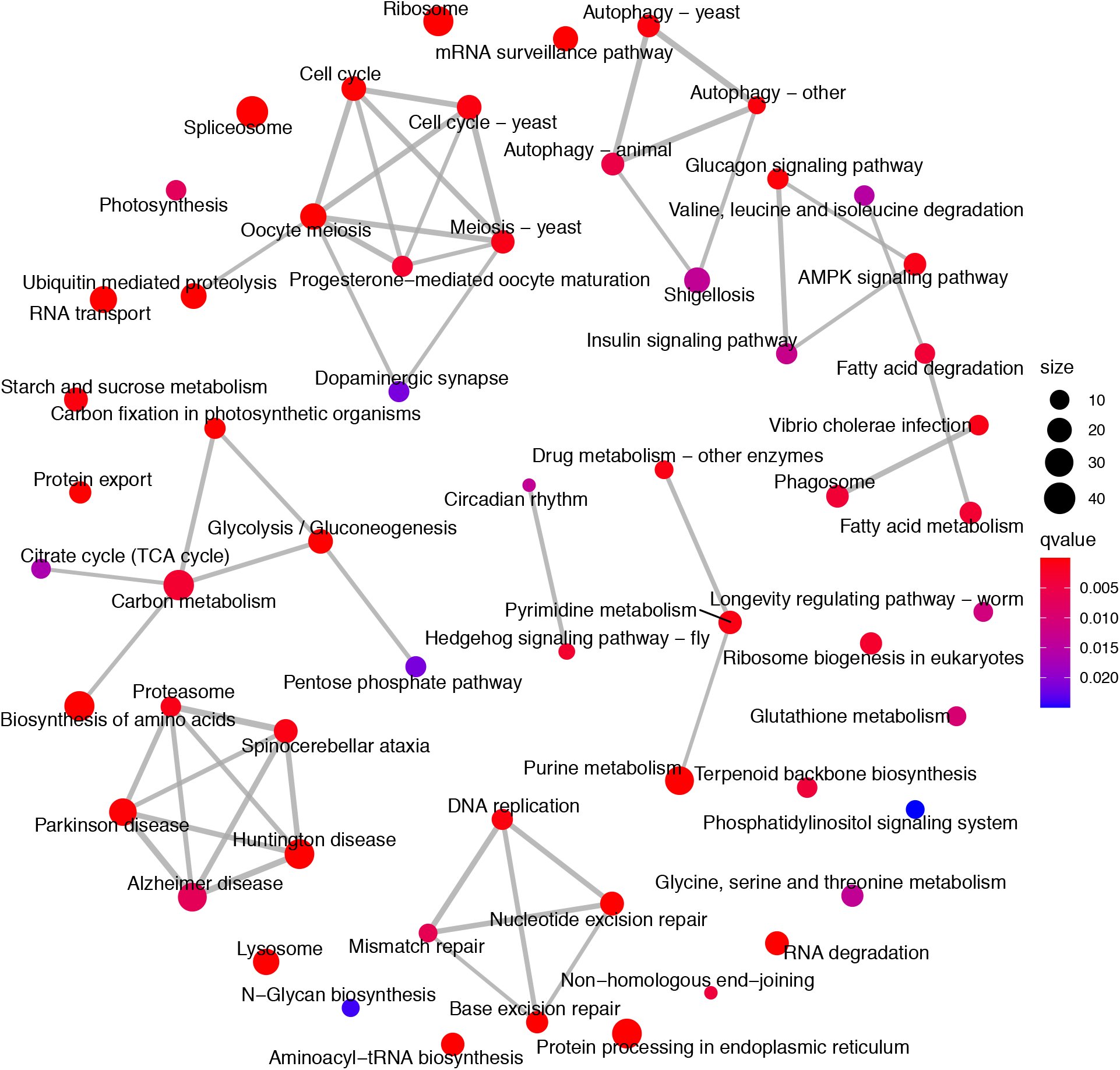
Enrichment map displaying the 58 significantly enriched KEGG pathways (circular nodes) resulting from KEGG enrichment analysis of the 11,514 differentially expressed transcripts between heat-stressed and control *Breviolum psygmophilum*. Gray lines connect pathways with overlapping genes. The color of each circle indicates the significance (q-value) and the size shows the number of genes in that pathway.

Because we measured multiple indices of photosynthetic performance that could assist with interpretation, we looked more closely at the two enriched pathways in our *B. psygmophilum* samples that were involved in photosynthesis. The enriched photosynthesis pathway included an ATP synthase subunit alpha (the regulatory subunit that facilitates or inhibits enzyme activity) that was significantly downregulated, with the largest log_2_ fold change in the pathway. Ferredoxin, which binds iron and sulfur and transfers electrons during photosynthesis, was also downregulated. Both the oxygen-evolving enhancer protein and P680 reaction center D1 proteins of photosystem II were significantly downregulated. Interestingly, the most upregulated gene was for the Psb27 protein involved in photosystem II assembly and repair. In addition, the two differentially expressed photosystem I reaction center subunit proteins were both upregulated, indicating shifting photosystem stoichiometry.

The pathway “Carbon fixation in photosynthetic organisms” was also enriched, and while multiple enzymes were differentially expressed, ribulose 1,5-bisphosphate carboxylase (RuBisCO) was not among them. The largest observed fold change corresponded to the significantly downregulated triose phosphate isomerase, which is involved in the regeneration phase of the reductive pentose phosphate cycle (Calvin-Benson Cycle). Other enzymes in this cycle were significantly downregulated, including a chloroplastic fructose-1,6-bisphosphatase, a fructose-bisphosphate aldolase, and a sedoheptulose-bisphosphatase. However, a ribulose-phosphate 3-epimerase was significantly upregulated, which catalyzes the conversion of xylulose 5-phosphate to ribulose 5-phosphate, and vice versa. Phosphoribulokinase, which converts ribulose 5-phosphate to ribulose 1,5-bisphosphate, was also upregulated. While ribulose-phosphate 3-epimerase is important for the regeneration phase of the reductive pentose phosphate cycle, it also plays a crucial role in the non-oxidative phase of the pentose phosphate pathway (which was also significantly enriched).

Many prior studies have pointed to the production of reactive oxygen species (ROS) as a trigger for symbiosis breakdown during thermal stress (Asada, 1996, McGinty *et al*., 2012, Venn *et al*., 2008, Weis, 2008). Thus, we examined enriched KEGG pathways that were involved in modulating oxidative stress, including the longevity regulating pathway and glutathione metabolism. In the longevity regulating pathway, both a glutathione S-transferase (which is also part of glutathione metabolism) and a superoxide dismutase were significantly upregulated. The superoxide dismutases are the first responders for detoxifying superoxide radicals, yielding oxygen and hydrogen peroxide. The isoform that was upregulated belongs to the family of superoxide dismutases that uses iron (Fe) or manganese (Mn) as a cofactor. Downregulated genes in this pathway included two that code for molecular chaperones involved in protein refolding and one that codes for a protease that degrades misfolded proteins.

Previously, the total glutathione content has been measured as an indicator of antioxidant capacity (Downs *et al*., 2002). The pathway for “Glutathione metabolism” was significantly enriched in *B. psygmophilum*, and included both a glutathione reductase and a glutathione peroxidase that were significantly downregulated. Glutathione reductase reduces glutathione-disulfide to produce glutathione in the chloroplast (Sies, 1999). Glutathione peroxidases, which protect cells from hydrogen peroxide, lipid peroxides, and organic hydroperoxide using glutathione, were all significantly downregulated. Contrastingly, glutathione S-transferases were significantly upregulated, which are important for scavenging ROS to maintain redox homeostasis.

In addition to the gene expression changes we found related to photosynthesis and oxidative stress, we found enriched pathways related to meiosis and the cell cycle. The “Oocyte meiosis” pathway included mostly downregulated genes. A protein involved in the structural maintenance of chromosomes and chromatid segregation was downregulated, and had the largest magnitude fold change. Separase, an enzyme implicated in meiotic chromosome separation, was also significantly downregulated. Likewise, in the enriched “Meiosis-yeast” pathway, the majority of differentially expressed genes were downregulated. Most of the differentially expressed genes in this pathway were also part of the “Oocyte meiosis” pathway. Notably, one meiotic recombination protein (*SPO11*) essential for genetic recombination during gamete production was significantly downregulated. Proteins involved in DNA replication were upregulated.

## Discussion

The symbiosis between corals and dinoflagellates in the family Symbiodiniaceae is often obligate in warm tropical, oligotrophic waters and heat stress can cause damage to the partners leading to dysbiosis. Because adult reef-building corals do not naturally exist in an aposymbiotic state, it is challenging to understand the role of either partner in the symbiosis under normal as well as under stressful conditions. The temperate coral *Astrangia poculata* provides an interesting system to understand the contribution of either partner to the symbiosis because adult healthy colonies exist aposymbiotically, and single colonies may contain both symbiotic and aposymbiotic polyps. Other coral hosts can be induced to be aposymbiotic, but this is often associated with significant stress to the host. While the facultatively symbiotic anemone Aiptasia is a useful model (Weis *et al*., 2008), this cnidarian does not build a skeleton and thus important cellular mechanisms differ between Aiptasia and tropical corals. Here we present data on how the cnidarian host responds to symbiosis under normal conditions and how that response changes when the symbiosis is challenged with heat stress. In this study, we characterized the heat stress response of paired symbiotic and aposymbiotic ramets of the temperate coral, *Astrangia poculata*, as well as its dominant algal symbiont *Breviolum psygmophilum*, using a combination of physiology measurements and gene expression analysis. Our study is among the first to describe a thermal stress response to elevated temperatures in *A. poculata* while controlling for coral host genotype, and the first to explore expression of both the host (*A. poculata*) and the symbiont (*B. psygmophilum*) during chronic high temperatures (∼3 weeks). Compared to the respective controls, heat-stressed symbiotic *A. poculata* ramets altered the expression of fewer host genes than heat-stressed aposymbiotic ramets as if the symbiotic polyp’s immune response was repressed by the algae. Additionally, more genes were differentially expressed between symbiotic and aposymbiotic controls than between heat-stressed symbiotic and aposymbiotic ramets. The application of heat stress minimized transcriptional differences between symbiotic and aposymbiotic ramets, indicating the symbiotic ramets were developing a bleached phenotype. However, the largest transcriptional response was observed in heat-stressed *B. psygmophilum*, which agreed with our measures of photosynthetic dysfunction. Taken together these data indicate that high temperature disrupts the function of the symbiont more than the host.

### Algal Photosynthesis was Impaired at High Temperatures

Examining trends in the maximum net photosynthesis rate *P*_max_ (when light is not limiting) shows how the capacity of symbiont photosynthesis changes with increasing temperatures. Over the course of the three-week experiment, the average *P*_max_ increased significantly from 18 to 22°C, but then remained relatively flat until 28°C. We then observed a significant decline in *P*_max_ above 28°C, indicating that the rate of inactivation of the photosynthetic proteins was greater than the rate of repair. The experimental treatment thus exceeded the thermal tolerance limit of the dinoflagellate symbiont, *Breviolum psygmophilum*. The reduction in *P*_max_ is likely caused by increased fluidity of lipids in the thylakoid membranes, yielding reduced capacity for electron transport (Iglesias-Prieto *et al*., 1992, Tchernov *et al*., 2004). While previous studies have measured photosynthesis rates with increasing temperatures and at multiple light intensities, this is the first study to construct photosynthesis-irradiance curves for *Breviolum psygmophilum* in hospite. One former paper showed a significant linear increase in photosynthesis rate between 15 and 27°C (Jacques *et al*., 1983), while a more recent study showed the change in photosynthesis rate over a larger temperature range between 6 and 32°C (Aichelman *et al*., 2019). A reduction in *P*_max_ demonstrates that less photosynthates are translocated to the host at high temperatures (Cantin *et al*., 2009, Jones *et al*., 1998) due to fewer symbionts remaining in host coral tissues and/or impaired photosynthetic capacity of the symbionts.

The sustained high values of maximum photochemical efficiency of photosystem II (Fv/Fm) from 18 to 27°C agreed with previous studies on *B. psygmophilum* in symbiosis and in culture (Burmester *et al*., 2017, Thornhill *et al*., 2008). However, above 27°C, the significant decrease in Fv/Fm indicated that the algal symbionts sustained damage to photosystem II. Photosystem II dysfunction is often measured as an indicator of tropical coral bleaching (Fitt *et al*., 2001, Warner *et al*., 1999). The reduced photosystem II activity could be the cause of decreased maximum photosynthesis rate in all heat-treated colonies by the end of the experiment. We did not see a change in photosynthetic efficiency (α, Figure 3B). A decrease in α would indicate that there was a reduction in the functional absorption cross section of photosystem II – a product of a reduction in the chlorophyll antenna size and/or number, and/or damage to chlorophyll reactions centers (Iglesias-Prieto & Trench, 1994). This means that the chlorophyll antenna was not necessarily damaged, and a decrease in the efficiency of light capture was not the cause of reduced maximum photosynthesis rate.

Instead, a reduction in the overall chlorophyll content could explain the decrease in reduced maximum net photosynthesis rate. A previous study showed that reflectance measurements are a good estimate for true chlorophyll absorption measurements (Enríquez *et al*., 2005) and we measured a significant decrease in the absorbance peak of chlorophyll *a* between initial (ambient temperature of 18°C) and final (30°C) conditions. This indicated that there was a decrease in the overall concentration of chlorophyll *a* pigments in the coral tissues that may explain the decrease in reduced maximum photosynthesis rate, although chlorophyll content could not be measured directly due to the limited availability of coral tissue.

### Respiration Response Curves Differed Between Symbiotic and Aposymbiotic Ramets

Our measures of respiration rates at increasing temperatures form the left side of a thermal performance curve, where the peak of the curve designates the point where performance is maximized (thermal optimum, *T*_opt_) (Huey & Stevenson, 1979, Schulte *et al*., 2011). Heat stress resulted in a reduction in maximum respiration rates between 29°C and 30°C in all of the symbiotic colonies while the aposymbiotic colonies of *A. poculata* showed variable responses in their respiration rates. The decrease in respiration observed in symbiotic colonies could have been a result of reduced algal respiration, reduced coral host respiration, or both. This difference indicates that the thermal performance curve for symbiotic colonies may be shifted relative to the curve for aposymbiotic colonies, with a *T*_opt_ around 29°C driven by the thermal tolerance of the symbiont. Because we did not see a uniform decrease in aposymbiotic coral respiration at the end of our experiment, we expect the *T*_opt_ to vary by coral host genotype, with some ramets potentially having a *T*_opt_ at 30°C or above.

Previous studies have compared the respiration rates of colonies of *A. poculata* with increasing temperatures. The authors of one study found little change in respiration rate between 11.5 and 23°C, before a significant increase in respiration between 23 and 27°C (Jacques *et al*., 1983). The highest temperature they used was 27°C, at which *A. poculata* colonies could survive indefinitely in the lab (Jacques *et al*., 1983). A more recent study measured changes in dark respiration rate for symbiotic and aposymbiotic colonies across a wide temperature range of 6 to 32°C. Interestingly, the latter study demonstrated differences in estimated thermal optima between colonies of *A. poculata* from different latitudes, although symbiotic and aposymbiotic colonies within each region did not have significantly different thermal optima (Aichelman *et al*., 2019).

These potential values for *T*_opt_ for Rhode Island *A. poculata* are higher than the maximum summer temperature of 23°C observed at our collection site (Burmester *et al*., 2017). However, *A. poculata* is capable of surviving in Rhode Island intertidal habitats (Grace, 2017), and thus may be adapted to much higher temperature extremes than those experienced regularly by subtidal colonies. Future studies are needed to fully measure the thermal response curves of *A. poculata* after reaching *T*_opt_. Had we continued to increase the water temperature for our experimental corals until we observed a steep drop in respiration rates corresponding to significant protein denaturation (Daniel & Danson, 2010, Sharpe & DeMichele, 1977), we would have risked significant mortality.

When the colonies were sampled for gene expression, the average respiration rate of the aposymbiotic colonies was increasing while symbiotic colonies were starting to decrease their respiration rates. At 30°C, algal cells had already sustained significant damage to photosystem II (Figure 3), and were likely translocating significantly less photosynthates to the coral host. This suggests that the coral alone has a higher thermal tolerance than the coral in symbiosis with *B. psygmophilum*, perhaps due to increased production of reactive oxygen species by the symbiont (Asada, 1996, McGinty *et al*., 2012, Venn *et al*., 2008, Weis, 2008). Interestingly, heat-stressed symbiotic colonies changed the expression of fewer genes than heat-stressed aposymbiotic colonies, relative to controls (see below).

### Symbiotic Ramets Altered the Expression of Fewer Genes than Aposymbiotic Ramets in Response to Heat Stress

When contrasting the heat stress response separately in symbiotic and aposymbiotic ramets of *A. poculata*, we found fewer differentially expressed genes between symbiotic control and symbiotic heat-stressed ramets (n=13) than we found between aposymbiotic control and heat-stressed ramets (n=76, Figure 5A). The overall low levels of differential expression align with recent work on aposymbiotic colonies of *A. poculata*, which demonstrated that seven times as many genes were differentially expressed under cold stress compared to heat stress in short-term experiments (Wuitchik *et al*., 2020). In response to chronic heat stress, aposymbiotic colonies in the present study mounted a significant inflammatory response. The most upregulated genes with informative annotation included a tumor necrosis factor-related protein, nitric oxide synthases, and a small heat shock protein. Tumor necrosis factor-related proteins activate proinflammatory reactions and regulate cell proliferation (Neumann *et al*., 2001). Nitric oxide synthases were highly upregulated in heat-stressed aposymbiotic colonies, and were also part of the enriched KEGG relaxin signaling pathway. The increased production of nitric oxide (NO) is part of the inflammation response, which is a protective mechanism for blocking injured or infected tissue from spreading (Ashley *et al*., 2012). Symbiotic sea anemones produced increased levels of NO in response to acute (24 hours) thermal stress, ultimately resulting in bleaching (Perez & Weis, 2006). In the same study, aposymbiotic anemones subjected to the same thermal stress did not increase NO production. The authors concluded that both symbiotic algae and host were required to increase NO production, yet this is in contrast with our experimental results with aposymbiotic *A. poculata*. In a later study, an immune elicitor was used to compare responses in symbiotic and aposymbiotic anemones. Interestingly, symbiotic animals produced significantly less NO than aposymbiotic animals, indicating an impaired immune response in symbiotic anemones (Detournay *et al*., 2012). Our results align with the observed reduced immune response in symbiotic animals.

We also observed a muted response to increased temperatures in our heat-stressed symbiotic colonies, relative to our control symbiotic colonies. These results indicate that symbiotic colonies of *A. poculata* have an overall weakened inflammatory response, enabling successful symbiosis establishment and maintenance of intracellular dinoflagellates. A general downregulation of the symbiotic cnidarian host’s inflammatory response under control conditions, relative to aposymbiotic hosts, was also found in a gene expression study of sea anemones (Lehnert *et al*., 2014). Other marine symbioses have displayed similar trends, including reduced NO signals in squid colonized by bioluminescent bacteria (Davidson *et al*., 2004). Future studies could measure NO production in symbiotic and aposymbiotic *A. poculata* directly using confocal microscopy. It would be informative to measure differences in this inflammatory response from acute as well as chronic heat stress to determine whether NO expression changes with the progression of the heat stress response.

Two other significantly enriched KEGG pathways implicated in the aposymbiotic coral heat stress response included the mitogen-activated protein kinase (MAPK) signaling pathway and the apoptosis pathway. The MAPK signaling cascade has been identified previously in cnidarian immunity, specifically related to wound healing (DuBuc *et al*., 2014). However, it is likely that components of MAPK signaling play important roles in multiple immune pathways (Mydlarz *et al*., 2016). In this study, we demonstrate a potential role for this pathway in the cnidarian heat stress response by increasing the expression of a molecular chaperone. Likewise, the apoptosis pathway is an important part of cnidarian immunity and is responsible for causing cell death to remove damaged cells (Mydlarz *et al*., 2016). The expression results for this pathway were indicative of apoptosis prevention under high temperatures. Altogether, the KEGG enrichment results point to an activated immune response in aposymbiotic ramets under chronic heat stress.

The most strongly downregulated genes with informative annotation in heat-stressed aposymbiotic colonies, relative to aposymbiotic controls, included enzymes involved in carbohydrate breakdown and fatty acid chain elongation. In addition, the KEGG pathway for protein digestion and absorption was significantly enriched and included four genes downregulated in heat-stressed aposymbiotic corals. This indicates that amino acid metabolism is also reduced, pointing to reduced catabolic processes under heat stress.

The symbiotic heat-stressed *A. poculata* colonies altered the expression of only 13 transcripts compared to controls. Three of the four genes that were upregulated were also upregulated in heat-stressed aposymbiotic colonies. The most upregulated gene coded for an influenza virus NS1A-binding protein homolog that was not differentially expressed in aposymbiotic colonies. This protein helps prevent cell death caused by destabilizing actin filaments (Matsudo *et al*., 2006). Thus, both symbiotic and aposymbiotic heat stressed colonies upregulated distinct genes involved in the negative regulation of apoptosis. However, all other immune response genes upregulated in aposymbiotic ramets were absent from the symbiotic *A. poculata* heat stress response. This indicates that symbiosis with *B. psygmophilum* alters the immune response of *A. poculata*.

### Symbiotic and Aposymbiotic Ramets Became More Transcriptionally Similar Under Heat Stress

Of the 125 differentially expressed transcripts between control symbiotic and aposymbiotic ramets, 123 of these were upregulated in symbiotic ramets. This demonstrates that additional cellular processes are necessary to maintain symbiosis with *B. psygmophilum*. Only 41 transcripts were still differentially expressed under heat stress conditions (Figure 5B). From a gene expression standpoint, the symbiotic and aposymbiotic ramets became more similar with the application of heat stress, indicating that the symbiotic ramets were developing an aposymbiotic phenotype.

KEGG enrichment analysis for the differentially expressed genes between symbiotic and aposymbiotic control ramets yielded multiple pathways involved in symbiosis maintenance. All genes involved in the pathways described in this section were more highly expressed in symbiotic corals. The lysosome pathway was particularly well annotated, and included the most genes of any pathway in this contrast. Lysosomes are a key component of proper immune function in marine invertebrates. Coral cells can rid the tissue of pathogens by phagocytosis as part of the innate immune response, wherein an invader is engulfed in a phagosome that fuses with a lysosome, and enzymes within the lysosome digest the unwanted material (Ellis *et al*., 2011). In a study of symbiosis establishment in sea anemone larvae, the lysosome pathway was similarly enriched and included four genes upregulated in symbiotic larvae (Wolfowicz *et al*., 2016). Increased activity of lysosomal peptidases and proteases is an associated response to bacterial invasion (Ellis *et al*., 2011). In addition, the autophagic machinery that includes digestion via lysosomes has been directly implicated in coral bleaching (Downs *et al*., 2009).

After coming in contact, the coral host must be able to distinguish the desired dinoflagellate symbiont from unwanted microbial invaders and food. Thus, the establishment and maintenance of the symbiosis could depend on cellular signaling mechanisms that are part of the antigen processing and presentation pathway, which was enriched in symbiotic corals under control conditions. The coral host phagocytic cell must be able to recognize the symbiotic dinoflagellate using specific cell receptors, and prevent the fusion of the endosome (symbiosome) containing the symbiont cell with lysosomes (Davy *et al*., 2012). Other than cathepsins, the only other protein included in the enriched antigen processing and presentation pathway was a gamma-interferon-inducible lysosomal thiol reductase (GILT), localized to the endosome in the reference KEGG pathway. This protein is critical for antigen processing and maintaining the efficiency of protein breakdown (Balce *et al*., 2014). Interestingly, the presence of reactive oxygen species alters the functioning of GILT, perhaps pointing to why this protein is no longer upregulated in heat-stressed symbiotic colonies.

Perhaps related to antigen processing was the enriched “other glycan degradation” pathway in control symbiotic colonies. Glycan-lectin interactions have been previously identified as important for host/symbiont recognition. The host membrane lectins bind glycans on the cell surface of Symbiodinaceae, initiating a cellular cascade leading to symbiosis establishment (Davy *et al*., 2012). Different Symbiodinaceae species differ in their glycomes, and thus the glycans of unwanted algal cells are likely broken down by the coral host (Logan *et al*., 2010).

Indeed the removal of glycans from the cell surfaces of Symbiodinaceae significantly reduced algal establishment in the aposymbiotic anemone *Aiptasia pulchella* (Lin *et al*., 2000).

Genes corresponding to the two co-functioning cholesterol transporters NPC1 and NPC2 were significantly upregulated in symbiotic colonies and were involved in both the enriched lysosome and cholesterol metabolism pathways. In a past study, NPC2-like proteins were upregulated in symbiotic anemones, relative to aposymbiotic anemones (Lehnert *et al*., 2014). Differences in cholesterol metabolism were expected, since cholesterol is an important component of cell membranes and is involved in cell signaling and intracellular transport. In addition, the cholesterol transporter NPC2 was implicated in cholesterol transport between an anemone host and symbiont (Oakley *et al*., 2016), consistent with our finding of upregulated NPC2 in symbiotic colonies of *A. poculata*. Because all four of these pathways are important for symbiosis establishment and maintenance, their reduced expression under increased temperatures indicated the onset of dysbiosis.

The 41 transcripts that were differentially expressed between symbiotic and aposymbiotic colonies, regardless of temperature, were all highly upregulated in symbiotic corals. The most highly upregulated genes included an arylsulfatase B, an amino acid transporter, an N-glycosylase/DNA lyase, and a mitochondrial ATP synthase subunit alpha. Arylsulfatase B is localized to lysosomes and is important for the regulation of cell adhesion and migration, and autophagy. Previously, autophagy was shown to be a potential mechanism for symbiont expulsion (bleaching), in conjunction with apoptosis (Dunn *et al*., 2007, Weis *et al*., 2008). N-glycosylase/DNA lyase is a DNA repair enzyme that binds damaged DNA in the nucleus. Combined thermal stress and increased solar radiation were shown to damage photochemistry in the algal symbiont while also leading to DNA damage in the host tissues of the Caribbean coral, *Orbicella* (=*Montastraea*) *faveolata* (Lesser & Farrell, 2004, Rodríguez-Román *et al*., 2006). Although, we only increased irradiance for short incubation periods to construct the photosynthesis-irradiance curves, we did observe reduced photochemical efficiency as well as a reduction in *P*_max_ for the symbiont. The combined impacts of high temperature stress and limited exposure to high irradiance could have resulted in DNA damage in *A. poculata*, necessitating the upregulation of a DNA repair enzyme.

Increased expression of a mitochondrial ATP synthase in symbiotic corals indicated an increased need for ATP in the coral cells. At the end of the heat stress experiment when corals were sampled for RNAseq, symbiotic colonies were reducing their respiration rates while aposymbiotic colonies had a more variable respiration response. The algal symbiont in the host tissue was clearly stressed at the end of the experiment, as indicated by both reduced photochemistry and reduced *P*_max_. The lost photosynthates from a damaged algal partner likely necessitated increased ATP production in the coral host. Contrastingly, a sea anemone decreased ATP synthase expression in response to acute thermal stress (Dunn *et al*., 2012), and a reef-building coral downregulated ATP synthase while bleached (Ricaurte *et al*., 2016). Heat exposure differed between these experiments and the current study, suggesting that the expression of ATP synthase varies depending on the duration and intensity of thermal stress.

### Breviolum psygmophilum responded to chronic heat stress by altering the expression of thousands of genes, including photosynthesis, antioxidant, and meiosis genes

Earlier studies found minimal transcriptional changes in Symbiodiniaceae strains under thermal stress (Barshis *et al*., 2014, Leggat *et al*., 2011, Putnam *et al*., 2013). However, more recently significant changes in Symbiodiniaceae intraspecies gene expression were described (Parkinson *et al*., 2016, Xiang *et al*., 2015), including relatively large transcriptional responses to heat stress resulting in 2,465 and 4,272 differentially expressed genes (Baumgarten *et al*., 2013, Levin *et al*., 2016). Here, we found 11,514 differentially expressed transcripts between control and heat-stressed samples of *Breviolum psygmophilum*, which corresponded to 58 significantly enriched KEGG pathways (Figure 6). It is possible that different species of Symbiodiniaceae respond to stress in fundamentally different ways. Future comparative studies with standardized heat stress exposures will be able to elucidate the underlying mechanisms behind these intriguing differences.

Two of the enriched KEGG pathways we identified – “Photosynthesis” and “Carbon fixation in photosynthetic organisms” were directly related to the reduction in photosynthetic activity that we measured in *B. psygmophilum*. In the photosynthesis pathway, the photosystem II oxygen-evolving enhancer protein (which regulates turnover of the photosystem II reaction center D1 protein) and the photosystem II P680 reaction center D1 protein were both downregulated, while two photosystem I subunit proteins were upregulated. This response is indicative of shifting stoichiometry between photosystems – part of the long-term photoacclimation response to counteract imbalances in excitation energy (Pfannschmidt *et al*., 1999, Pfannschmidt *et al*., 2001). In addition, reduced expression of both ATP synthase and ferredoxin indicate reduced capacity for electron transport and repressed photosynthesis. These expression results agree with the observed reduction in *P*_max_ in heat stressed *B. psygmophilum*. Because the ATP generated from electron transport is needed to power carbon fixation, the excess light energy needs to be dissipated through other pathways, such as through the formation of reactive oxygen species (Roth, 2014).

Notably, the key enzyme RuBisCO in the “Carbon fixation in photosynthetic organisms” pathway was not differentially expressed between heat stressed and control *B. psygmophilum*. Instead, the downregulation of multiple genes involved in this KEGG pathway and the upregulation of the enzyme ribulose-phosphate 3-epimerase that is also involved in the pentose phosphate pathway indicated a shift from carbon fixation to glucose catabolism. The “Pentose Phosphate Pathway”, which was also significantly enriched, yields reducing equivalents (NADPH) that protect cells from oxidative stress (Liang *et al*., 2011). The addition of hydrogen peroxide to intact chloroplasts significantly inhibits CO_2_ fixation (Kaiser, 1976, Kaiser, 1979) while activating the oxidative pentose phosphate pathway (Kaiser, 1979). Therefore, elevated levels of reactive oxygen species in *B. psygmophilum* cells could be causing the observed changes in the expression of genes involved in carbon fixation and the pentose phosphate pathway.

The differentially expressed genes associated with the enriched longevity regulating and glutathione metabolism pathways further support that heat-stressed *B. psygmophilum* were experiencing oxidative stress at the final temperature of 30°C. *B. psygmophilum* subjected to chronic high temperatures upregulated an isoform of superoxide dismutase in the Fe-Mn family involved in detoxifying superoxide radicals (Bowler *et al*., 1991). Multiple isoforms of MnSOD have been described in Symbiodiniaceae species previously (Krueger *et al*., 2015), with potentially differing subcellular locations. Prior work has shown that different Symbiodiniaceae species produce ROS at elevated temperatures (McGinty *et al*., 2012), and some respond to thermal stress by increasing superoxide dismutase expression in culture (Gierz *et al*., 2017, Goyen *et al*., 2017). Further, a glutathione S-transferase in both the longevity regulating and glutathione metabolism pathways was significantly upregulated. Glutathione S-transferase is involved in removing peroxides and electrophiles that could damage DNA, RNA, and proteins (Noctor *et al*., 2002). High temperatures have been shown to increase glutathione S-transferase activity in addition to superoxide dismutase activity in a congeneric Symbiodiniaceae species in culture, providing additional antioxidant capabilities (Krueger *et al*., 2014). The current study builds on these results and demonstrates that upregulating superoxide dismutase and glutathione S-transferase is part of the *B. psygmophilum* heat stress response in hospite.

The potential impacts of temperature stress on sexual reproduction in *B. psygmophilum* were compelling, given our limited understanding of drivers of sex in the Symbiodiniaceae. The enriched “Oocyte meiosis” and “Meiosis – yeast” pathways contained multiple meiosis-specific genes that were significantly downregulated. One of these genes coded for the meiotic recombination protein SPO11, which is necessary for creating breaks in double-stranded DNA (Romanienko & Camerini-Otero, 2000), thereby enabling meiotic recombination. SPO11 also plays a role in proper meiotic synapsis, or the pairing of homologous chromosomes in prophase I (Romanienko & Camerini-Otero, 2000). The downregulation of multiple meiosis-related genes in *B. psygmophilum* under high temperatures indicated suppressed sexual reproduction. While meiosis-specific genes have been described in Symbiodiniaceae genomes (Chi *et al*., 2014), cytological evidence is lacking. A prior study examining the heat stress response of two populations of cultured *Cladocopium* sp. (formerly type C1) found significant upregulation of meiosis-specific genes, including SPO11 (Levin *et al*., 2016). The contrasting patterns of meiosis gene expression in *B. psygmophilum* (this study) and *Cladocopium* spp. (Levin *et al*., 2016) after over a week of increased temperatures could indicate different strategies for regulating gamete production in response to stress between the species, or differences between cultured cells and those in hospite. Further studies of when, where and how sexual reproduction occurs in the Symbiodiniacae are needed.

## Conclusion

The results from this experiment indicate that *B. psygmophilum* mounts a significant transcriptional response to chronic high temperature stress by altering the expression of genes involved in photosynthetic dysfunction as well as mediating oxidative stress. In contrast, symbiotic colonies of *A. poculata* altered the expression of very few host genes compared to aposymbiotic colonies, which activated an inflammatory response under high temperatures. This suggests that *B. psygmophilum* suppresses gene expression in its coral host, perhaps by inhibiting aspects of the inflammatory response of the coral (Davidson *et al*., 2004, Lehnert *et al*., 2014). Specifically, it appears that the presence of the symbiont results in suppressed nitric oxide synthase expression at chronic high temperatures. As a result, thermally stressed symbiotic colonies of *A. poculata* may be more susceptible to pathogens (Bogdan, 2001, Chakravortty & Hensel, 2003). Future work on symbiotic and aposymbiotic *A. poculata* could investigate whether a differential response to harmful microbes exists under normal and high temperature conditions. A growing body of work is demonstrating the link between global warming and increased disease prevalence in reef-building corals (Bruno *et al*., 2007, Harvell *et al*., 1999, Precht *et al*., 2016, Randall & Van Woesik, 2015, Ruiz-Moreno *et al*., 2012). The *A. poculata*-*B. psygmophilum* system may prove to be a tractable model for understanding the interaction between chronic thermal stress and immune challenge in scleractinian corals.

## Acknowledgements

This work would not have been possible without the support of excellent undergraduate students, including Andrew Samulewicz, Kyle Trabocco, Timothy Gilpatrick, and Tanner Quiggle who helped build and maintain aquarium systems and care for experimental animals. Sheila Kitchen and Viridiana Avila provided guidance and advice on molecular techniques and the differential expression analysis. Funding for this research was provided by a Penn State Institutes of Energy and the Environment Seed Grant awarded to I.B.B. A.N.C. was supported by the National Science Foundation (NSF) Graduate Research Fellowship Program under Grant No. DGE1255832. The conclusions are those of the authors and do not necessarily reflect the views of the NSF.

## Data Availability

Sequences will be deposited at the NCBI Sequence Read Archive, and supplementary tables of gene annotations, figures, and raw physiology data will be made available on the Penn State ScholarSphere institutional repository upon publication.

## Author Contributions

ANC, LAG, RIP, and IBB designed the experiment. ANC and LAG conducted the experiment, and analyzed physiology data. ANC completed all molecular work and differential expression analyses, and wrote the paper. JRF and EMB assembled the reference coral host transcriptome. EMB and RDR collected the coral colonies used in the experiment. IBB and RIP provided funding and materials. All authors read and approved the final manuscript.

## Notes

### Competing Interest Statement

The authors have declared no competing interest.

